# Design principles for robust vesiculation in clathrin-mediated endocytosis

**DOI:** 10.1101/050484

**Authors:** Julian E. Hassinger, George Oster, David G. Drubin, Padmini Rangamani

## Abstract

A critical step in cellular trafficking pathways is the budding of membranes by protein coats, which recent experiments have demonstrated can be inhibited by elevated membrane tension. The robustness of processes like clathrin-mediated endocytosis (CME) across a diverse range of organisms and mechanical environments suggests that the protein machinery in this process has evolved to take advantage of some set of physical design principles to ensure robust vesiculation against opposing forces like membrane tension. Using a theoretical model for membrane mechanics and membrane protein interaction, we have systematically investigated the influence of membrane rigidity, curvature induced by the protein coat, area covered by the protein coat, membrane tension and force from actin polymerization on bud formation. Under low tension, the membrane smoothly evolves from a flat to budded morphology as the coat area or spontaneous curvature increases, whereas the membrane remains essentially flat at high tensions. At intermediate, physiologically relevant, tensions, the membrane undergoes a *snapthrough instability* in which small changes in the coat area, spontaneous curvature or membrane tension cause the membrane to “snap” from an open, U-shape to a closed bud. This instability can be smoothed out by increasing the bending rigidity of the coat, allowing for successful budding at higher membrane tensions. Additionally, applied force from actin polymerization can bypass the instability by inducing a smooth transition from an open to a closed bud. Finally, a combination of increased coat rigidity and force from actin polymerization enables robust vesiculation even at high membrane tensions.

**Significance statement:** Plasma membrane tension plays an important role in various biological processes. In particular, recent experimental studies have shown that membrane tension inhibits membrane budding processes like clathrin-mediated endocytosis (CME). We have identified a mathematical relationship between the curvature-generating capability of the protein coat and membrane tension that can predict whether the coat alone is sufficient to produce closed buds. Additionally, we show that a combination of increased coat rigidity and applied force from actin polymerization can produce closed buds at high membrane tensions. These findings are general to any membrane budding process, suggesting that biology has evolved to take advantage of a set of physical design principles to ensure robust vesicle formation across a range of organisms and mechanical environments.

**Author Contributions:** J.E.H., G.O., and P.R. designed research. J.E.H. performed research. J.E.H., D.G.D., and P.R. analyzed data. J.E.H., G.O., D.G.D., and P.R. wrote the paper.

## Introduction

Clathrin-mediated endocytosis (CME), an essential cellular process in eukaryotes, is an archetypical example of a membrane deformation process that takes as input multiple variables such as membrane bending, tension, protein-induced spontaneous curvature and actin-mediated forces, and generates vesicular morphologies as its output [1]. Though more than 60 different protein species act in a coordinated manner during CME [2], we can distill this process into a series of mechanochemical events where a feedback between the biochemistry of the protein machinery and the mechanics of the plasma membrane and the actin cytoskeleton control endocytic patch topology and morphology [3, 4].

In Figure 1, we outline the main steps that lead to bud formation. Despite the complexity of CME, a variety of experimental approaches have served to identify the governing principles of bud formation in CME. We have identified a few key features from recent experiments that govern bud formation and have summarized the main results below.

**Figure 1.**
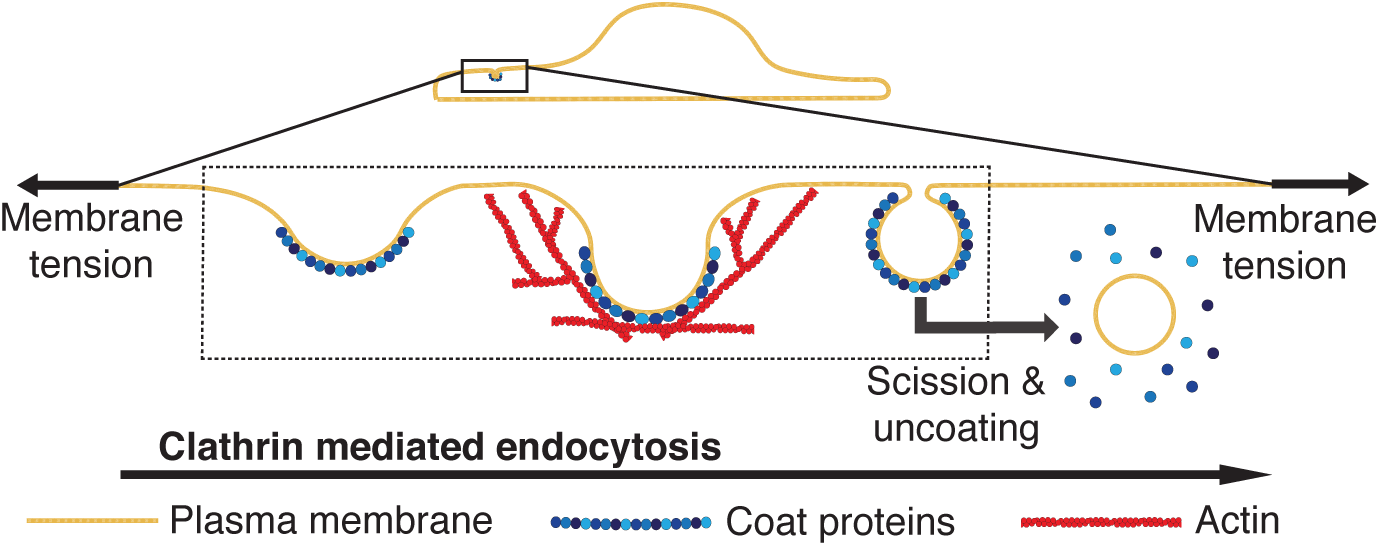
Schematic depiction of the main mechanical steps in clathrin-mediated endocytosis. A multi-component protein coat forms on the plasma membrane and causes the membrane to bend inwards, forming a shallow pit. As the coat matures, the membrane becomes deeply invaginated to form an open, U-shaped pit before constricting to form a closed, Ω-shaped bud. The bud subsequently undergoes scission to form an internalized vesicle and the coat is recycled. Actin polymerization is thought to provide a force, **f**, to facilitate these morphological changes, particularly at high membrane tensions [5]. Our study is focused on understanding the impact of membrane tension on the morphological changes effected by the coat and actin polymerization, as indicated by the dashed box.

- **Protein-induced spontaneous curvature:** A critical step in CME is the assembly of a multicomponent protein coat that clusters cargo and bends the membrane into a budded morphology. Clathrin assembles into a lattice-like cage on the membrane with the assistance of adaptor proteins that directly bind lipids [6, 7]. This assembly is generally thought to act as a scaffold that imposes its curvature on the underlying membrane [8]. Recent work suggests that other components of the coat can also contribute to membrane bending through scaffolding by F-BAR domains, amphipathic helix insertion into the bilayer, and adaptor protein crowding [6, 9–11]. Crowding of cargo molecules on the outer leaflet of the plasma membrane opposes invagination of the membrane [11, 12]; we can think of this effect as simply a negative contribution to the curvature of the coat. The contributions from each of these membrane bending mechanisms can be combined into a single measure of the curvature generating capability of the coat, or spontaneous curvature, with an effective strength that depends on its composition, density and area coverage [13, 14].
- **Membrane properties (moduli):** The bending modulus, or rigidity, of the plasma membrane is a material property of the lipid bilayer describing its resistance to bending, and is determined by its composition [15]. This bending rigidity is generally thought to be the primary opposing force to membrane deformations [16]. Supporting this idea, a decrease in the bending rigidity of the plasma membrane by incorporation of polyunsaturated phospholipids was found to stimulate an uptake of transferrin, a hallmark of increased endocytic dynamics [17].
- **Membrane tension:** The plasma membrane of animal cells is under tension as a result of in-plane stresses in the bilayer and connections between the membrane and the underlying actomyosin cortex [18, 19]. It has been demonstrated *in vitro* that membrane tension opposes deformations to the membrane by curvature-generating proteins [20]. *In vivo*, elevated tension in combination with actin inhibitors causes clathrin-coated pits (CCPs) to exhibit longer lifetimes and increased the number of long-lived, presumably stalled, pits [5]. Under these conditions, open, U-shaped pits were found to be enriched as compared to closed, Ω-shaped pits when visualized by electron microscopy [5, 21]. Similar observations have been made in a reconstituted system where purified coat proteins were able to substantially deform synthetic lipid vesicles under low tension but were stalled at shallow, U-shaped pits at a higher tension [22]. Additionally, membrane tension has been shown to induce disassembly of caveolae [23] as well as flattening of exocytic vesicles following fusion to the plasma membrane [24].
- **Force from actin polymerization:** It has long been appreciated that actin polymerization is an essential component in the CME pathway in yeast [25], presumably due to the high turgor pressure in this organism [26, 27]. In recent years, it has become clear that actin plays an important role in mammalian CME in conditions of high membrane tension [5] and to uptake large cargos like virus particles [28, 29].

From these studies, we can conclude that there are multiple variables that control the budding process, and are particularly dependent on the cell type and specific process. In whole cells, many different variables are at play simultaneously. There remain substantial challenges associated with identifying the separate contributions from each of these factors through experimental approaches. The diffraction-limited size of CCPs (~ 100 nm) makes it currently impossible to directly image the morphology of the membrane in situ in living cells. The temporal regularity of yeast CME has allowed for the visualization of time-resolved membrane shapes in this organism using correlative fluorescence and electron microscopy [30, 31]. However, this approach is quite difficult to use in mammalian cells because of the wide distribution of CCP lifetimes [32, 33]. Additionally, current techniques are only capable of measuring global tension [19, 34, 35], making it nearly impossible to determine how local membrane tension impacts the progression of membrane deformation at a given CCP. Beyond this, it is difficult to perturb the composition and tension of the plasma membrane in a controlled and quantitative way.

Reconstitution of membrane budding *in vitro* allows for control of lipid and protein composition as well as membrane tension [8, 22, 36]. However, coat area is an uncontrolled variable in these studies, and explicitly varying the spontaneous curvature would be challenging because the connection between individual molecular mechanisms of curvature generation and spontaneous curvature is not fully understood. Additionally, controlled application of force from actin polymerization at single sites of membrane budding has not yet been possible.

For these reasons, we have chosen to pursue a computational approach that allows us to explore how each of the factors that governs budding contributes to morphological progression of membrane budding, when varied in isolation or in various combinations.

Mathematical modeling has proven to be a powerful approach to describe observed shapes of membranes in a wide variety of contexts, from shapes of red blood cells to shape transformations of vesicles [13, 37]. In recent years, mathematical modeling has provided insight into various aspects of membrane deformation in number of budding phenomena including domain-induced budding, caveolae, ESCRTs, and CME [38–40]. For example, Liu et al. showed that a line tension at a lipid phase boundary could drive scission in yeast [3, 41], while Walani et al. showed that scission could be achieved via snapthrough transition at high membrane tension [42]. These studies and others [27, 43, 44] have demonstrated the utility of membrane modeling approaches for studying CME. However, none have systematically explored how the various parameters described above come together to determine the success or failure of budding.

In this study, we seek to answer the following questions: How does membrane tension affect the morphological progression of endocytic pits? How do the various mechanisms of membrane bending interact to overcome the effects of high tension and form buds? What are the design principles for robust vesiculation?

## Model development

### Membrane mechanics

We model the lipid bilayer as a thin elastic shell. The bending energy of the membrane is modeled using the Helfrich-Canham energy, which is valid for radii of curvatures much larger than the thickness of the bilayer [13]. Since the radius of curvature of typical endocytic patch is ≈ 50 nm [45, 46], application of this model provides a valid representation of the shapes of the membrane. Further, we assume that the membrane is at mechanical equilibrium at all times. This assumption is reasonable because clathrin-mediated endocytosis occurs over a timescale of tens of seconds [2, 5, 32, 33], and the membrane has sufficient time to attain mechanical equilibrium at each stage [3, 27]. We also assume that the membrane is incompressible/inextensible because the energetic cost of stretching the membrane is high [47]. This constraint is implemented using a Lagrange multiplier (see SI Appendix, Section 1 for details). Finally, for simplicity in the numerical simulations, we assume that the endocytic patch is rotationally symmetric (SI Appendix, Figure S1).

### Membrane-protein interaction: spontaneous curvature and area of coat

One of the key features of CME is coat-protein association with the plasma membrane. We model the strength of curvature induced by the coat proteins with a spontaneous curvature term (C). Spontaneous curvature represents an asymmetry (e.g. lipid composition, protein binding, shape of embedded proteins) across the leaflets of the membrane that favors bending in one direction over the other with a magnitude equal to the inverse of the preferred radius of curvature [13]. In our case, the spontaneous curvature represents the preferred curvature of the coat proteins bound to the cytosolic face of the membrane, consistent with its usage in other studies [20, 27, 42, 48, 49].

Our model reflects the fact that the clathrin coat covers a finite area and that this region has different physical properties (e.g. spontaneous curvature, bending rigidity) than the surrounding uncoated membrane. Heterogeneity in the spontaneous curvature and bending rigidity is accommodated by using a local rather than global area incompressibility constraint [50-52]. This allows us to simulate a clathrin coat by tuning the area, spontaneous curvature, and rigidity of the coated region with respect to the uncoated membrane.

### Governing equations

We use a modified version of the Helfrich energy that includes spatially-varying spontaneous curvature *C* (*θ^α^*) bending modulus *κ*(*θ^α^*), and Gaussian modulus *κ_G_*(*θ^α^*) [42, 48, 50, 52],

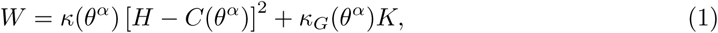

where *W* is the energy per unit area, *H* is the local mean curvature, and *K* is the local Gaussian curvature. *θ^α^* denotes the surface coordinates where *α* ∊ {1,2}. This form of the energy density accommodates the coordinate dependence or local heterogeneity in the bending modulus *κ*, Gaussian modulus *κ_G_,* and the spontaneous curvature *C*, allowing us to study how the local variation in these properties will affect budding. Note that this energy functional differs from the standard Helfrich energy by a factor of 2, with the net effect being that our value for the bending modulus, *κ*, is twice that of the standard bending modulus typically encountered in the literature.

A balance of forces normal to the membrane yields the “shape equation” for this energy functional,

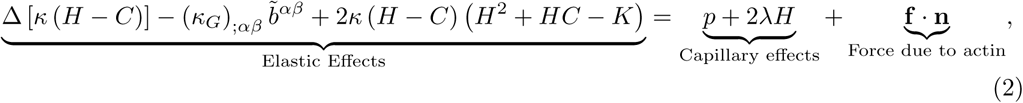

where ∆ is the surface Laplacian, *p* is the pressure difference across the membrane, λ is interpreted to be the membrane tension, *b^αβ^* are components of the curvature tensor, **f** is a force per unit area applied to the membrane surface, and **n** is the unit normal to the surface [42, 50]. In this model, **f** represents the applied force exerted by the actin cytoskeleton; this force need not necessarily be normal to the membrane. In this work, the transmembrane pressure is taken to be *p* = 0 to focus on the effect of membrane tension.

A consequence of heterogenous protein-induced spontaneous curvature, heterogeneous moduli, and externally applied force is that λ is not homogeneous in the membrane [48, 50]. A balance of forces tangent to the membrane yields the spatial variation of membrane tension,

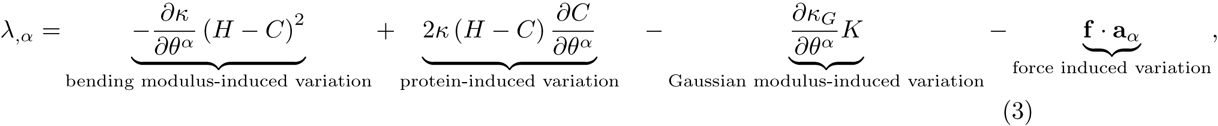

where (·) _,*α*_ is the partial derivative with respect to the coordinate *α* and **a**_*α*_ is the unit tangent in the *α* direction. λ can be interpreted as the membrane tension [48, 52], and is affected by the spatial variation in spontaneous curvature and by affected by the tangential components (**a**_*α*_) of the force due to the actin cytoskeleton. A complete derivation of the stress balance and the governing equations of motion is presented in SI Appendix, Section 1.

## Results

### Membrane tension controls bud formation by curvature-generating coats

In order to understand how membrane tension affects the morphology of a coated membrane, we performed two sets of calculations. In the first set, we studied the effect of varying coat area and membrane tension on membrane budding in the absence of external forces from the actin network. Simulations were performed by increasing the area of a curvature-generating coat at the center of an initially flat patch of membrane. We maintained the spontaneous curvature of the coat to be constant as *C*_0_ = 0.02 nm^−1^ in the coated region with a sharp transition at the boundary between the coated and bare membrane (implemented via hyperbolic tangent functions, SI Appendix, Figure S2). The membrane tension was varied by setting the value of λ at the boundary of the membrane patch, which corresponds to the tension in the surrounding membrane reservoir.

High membrane tension (0.2 pN/nm) inhibits deformation of the membrane by the protein coat (Figure 2A, upper row). As the area of the coated region (*A*_coat_) increases, the membrane remains nearly flat, and the size of the coated region can grow arbitrarily large without any substantial deformation (SI Appendix, Figure S3; Animation A1, left). The spontaneous curvature of the coat is simply unable to overcome the additional resistance provided by the high membrane tension. In contrast, at low membrane tension (0.002 pN/nm), increasing the coat area causes a smooth evolution from a shallow to deep U-shape to a closed, Ω-shaped bud. (Figure 2A, lower row; Animation A1, right). We stopped the simulations when the membrane is within 5 nm of touching at the neck, at which point bilayer fusion resulting in vesicle scission is predicted to occur spontaneously [41, 53]. These morphological changes are similar to those observed in clathrin-mediated endocytosis [46] and do not depend on the size of the membrane patch (SI Appendix, Figure S4).

**Figure 2.**
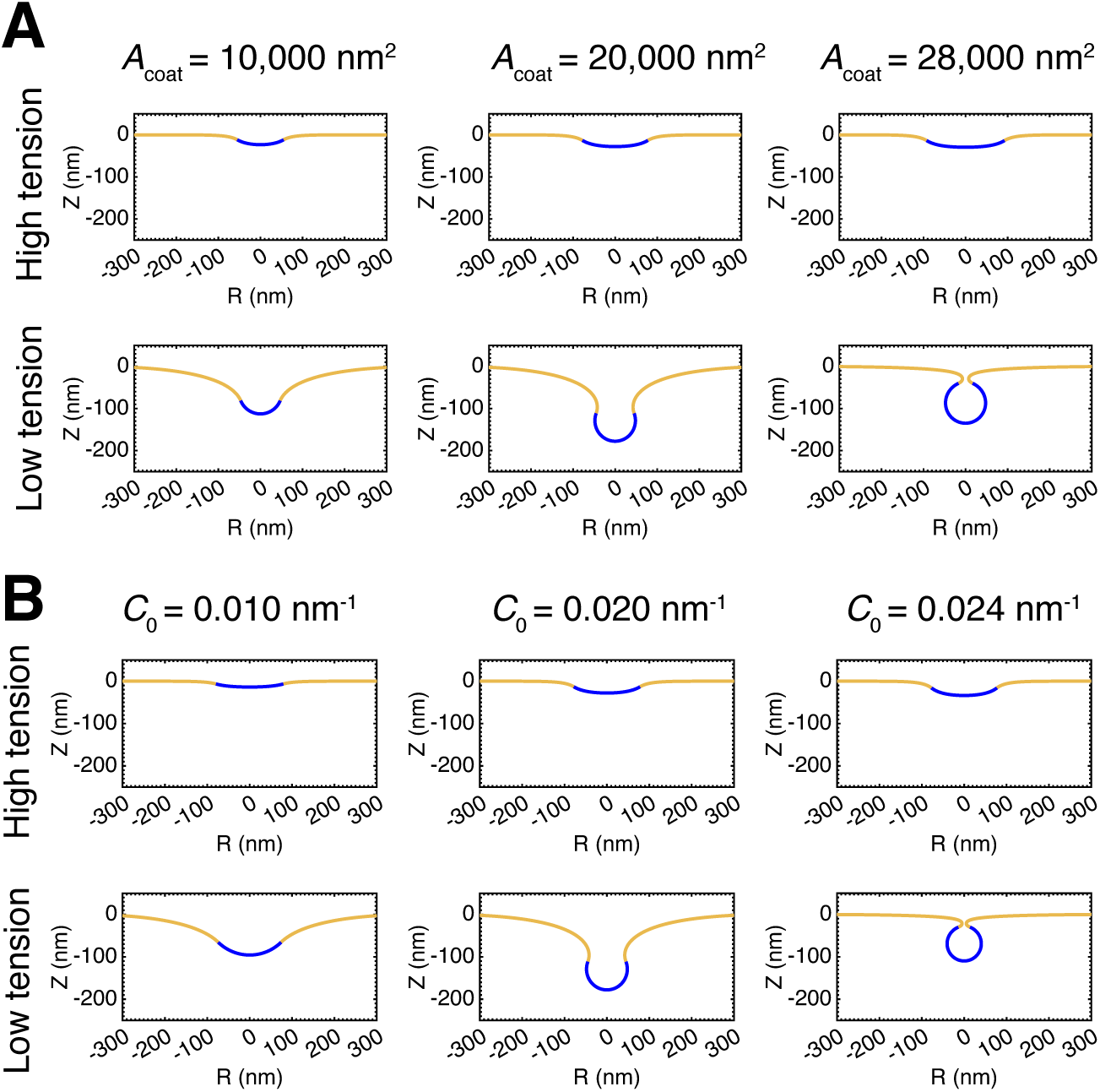
Membrane tension inhibits the ability of curvature generating coats to induce budding. **(A)** Profile views of membrane morphologies generated by simulations in which the area of a curvature-generating coat progressively increases, covering more of the bare membrane. The curvature-generating capability, or spontaneous curvature, of the coat is set at *C*_0_ = 0.02 nm^−1^, corresponding to a preferred radius of curvature of 50 nm [12]. *Upper row*: High membrane tension, λ_0_ = 0.2pN/nm. The membrane remains nearly flat as the area of the coat increases. *Lower row*: Low membrane tension, λ_0_ = 0.002 pN/nm. Addition of coat produces a smooth evolution from a flat membrane to a closed bud. **(B)** Membrane profiles for simulations with a constant coat area in which the spontaneous curvature of the coat progressively increases. The area of the coat is *A*_coat_ = 20, 106 nm^2^. *Upper row:* High membrane tension, λ_0_ = 0.2pN/nm. The membrane remains nearly flat with increasing spontaneous curvature. *Lower row:* Low membrane tension, λ_0_ = 0.002 pN/nm. Increasing the spontaneous curvature of the coat induces a smooth evolution from a flat membrane to a closed bud.

Since increasing coat area alone could not overcome the tension effects of the membrane, we asked if increasing the spontaneous curvature of the coat overcomes tension-mediated resistance to deformation. To answer this question, we performed simulations in which the spontaneous curvature of the coat increases while the area covered by the coat remains constant at approximately the surface area of a typical clathrin coated vesicle, *A*_coat_ = 20, 106 nm^2^[46]. As before, high membrane tension (Figure 2B, upper row; Animation A2, left) prevents deformation of the membrane by the coat. Even increasing the spontaneous curvature to a value of 0.04 nm^−1^, corresponding to a preferred radius of curvature of 25 nm and twice the value used the coat growing simulations, does not produce a closed bud (SI Appendix, Figure S5). In the case of low membrane tension (Figure 2B, lower row; Animation A2, right), a progressive increase in the coat spontaneous curvature causes a smooth evolution from a shallow to deep U-shape to a closed, Ω-shaped bud. The similarity between the membrane morphologies in Figure 2A and 2B indicates that the interplay between spontaneous curvature, coat area, and membrane tension is a governs membrane budding.

### Transition from U- to Ω-shaped buds occurs via instability at intermediate membrane tensions

Experimentally measured membrane tensions in mammalian cells typically fall between the high and low tension regimes presented in Figure 2 [54]. At an intermediate, physiologically relevant value of membrane tension (0.02 pN/nm), increasing the area of the coat causes substantial deformation of the membrane (Figure 3A). However, the transition from an open to a closed bud is no longer smooth. Figure 3A shows a bud just before (dashed line) and after (solid line) a small amount of area is added to the coat. This small change in area causes the bud to “snap” closed to an Ω-shaped morphology (Animation A3). This situation is known as a *snapthrough instability*, and similar instabilities have been observed in other recent membrane modeling studies [27, 42]. We emphasize that these are two equilibrium shapes of the membrane, and the exact dynamical transition between these states (i.e. intermediate unstable shapes and timescale) is not modeled here.

To visualize why this abrupt transition should occur, Figure 3B plots the mean curvature at the tip of the bud as a function of the area of the coat. In comparison to the high and low membrane tension cases (SI Appendix, Figure S6), there are two branches of equilibrium shapes of the membrane. The lower and upper branches represent “open” and “closed” morphologies of the bud, respectively. The marked solutions indicate the two morphologies depicted in Figure 3A. The open bud in Figure 3A is at the end of the open bud solution branch, so any addition of small area to the coat necessitates that the membrane adopt a closed morphology

**Figure 3.**
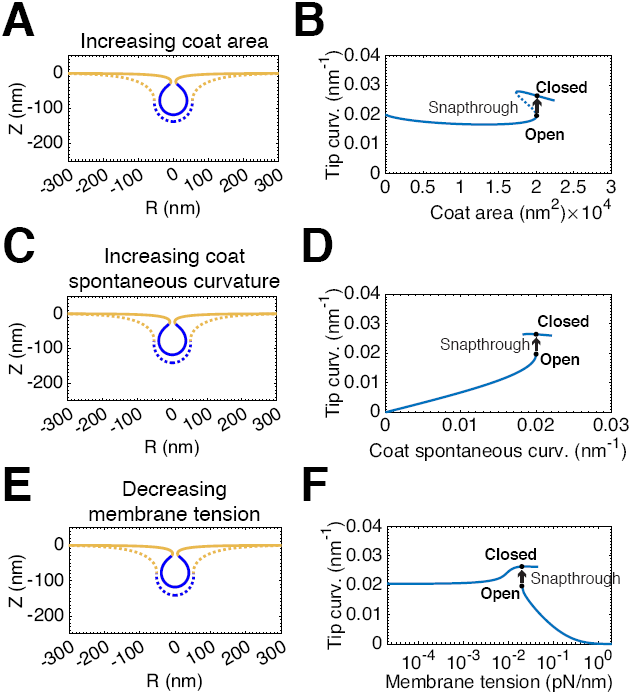
A snapthrough instability exists at intermediate, physiologically relevant [54], membrane tensions, λ_0_ = 0.02pN/nm. **(A)** Membrane profiles showing bud morphology before (dashed line, *A*_coat_ = 20,065 nm^2^) and after (solid line, *A*_coat_ = 20,105 nm^2^) addition of a small amount of area to the coat, C_0_ = 0.02 nm^−1^. **(B)** Mean curvature at the tip of the bud as a function of the coat area. There are two stable branches of solutions of the equilibrium membrane shape equations. The lower branch consists of open, U-shaped buds while the upper branch consists of closed, Ω-shaped buds. The dashed portion of the curve indicates “unstable” solutions that are not accessible by simply increasing and decreasing the area of the coat. The marked positions on the curve denote the membrane profiles shown in (A). The transition between these two shapes is a snapthrough instability, in which the bud “snaps” closed upon a small addition to area of the coat. **(C)** Bud morphologies before (dashed line) and after (solid line) a snapthrough instability with increasing spontaneous curvature, *A*_coat_ = 20, 106 nm^2^, *C*_0_ = 0.02nm^2^. **(D)** Mean curvature at the tip of the bud as a function of the spontaneous curvature of the coat. **(E)** Bud morphology before (dashed line) and after (solid line) a snapthrough instability with decreasing membrane tension, *A*_coat_ = 20, 106 nm^2^, *C*_0_ = 0.02nm^2^, λ_0_ = 0.02pN/nm. **(F)** Mean curvature at the tip of the bud as a function of the membrane tension.

This instability is also present for situations with increasing coat spontaneous curvature and constant coat area (Animation A4). Figure 3C shows membrane profiles before (dashed line) and after (solid line) a snapthrough transition triggered by an increase in spontaneous curvature. Figure 3D plots the mean curvature at the bud tip as a function of the coat spontaneous curvature. Similarly to Figure 3B, we observe that there are two branches of equilibrium membrane shapes.

Additionally, this instability is encountered when membrane tension is varied and the coat area and spontaneous curvature are maintained constant (Animation A5). Figure 3E shows membrane profiles before (dashed line) and after (solid line) a snapthrough transition triggered by a decrease in membrane tension. In Figure 3F we again see two solution branches in the plot of mean curvature at the tip as a function of membrane tension indicating a discontinuous transition between open and closed buds as tension is varied.

### The instability exists over a range of membrane tensions, coat areas, and spontaneous curvatures

Over what ranges of tension and spontaneous curvature does this snapthrough transition occur? First, to understand the nature of the transition between low and high membrane tension regimes, we performed simulations over several orders of magnitude of the membrane tension (10^−1^ to 1 pN/nm), encompassing the entire range of measured physiological tensions [54], as well as over a range of spontaneous curvatures of the coat (0 to 0.05 nm^−1^), corresponding to preferred radii of curvature from 20 nm and up. Based on the results, we constructed a phase diagram summarizing the observed morphologies (Figure 4A). The blue region denotes a smooth evolution to a closed bud, the red region represents a failure to form a closed bud, and green region indicates a snapthrough transition from an open to a closed bud. This phase diagram clearly shows that the distinction between “low” and “high” membrane tension conditions depends on the magnitude of the spontaneous curvature of the coat.

**Figure 4.**
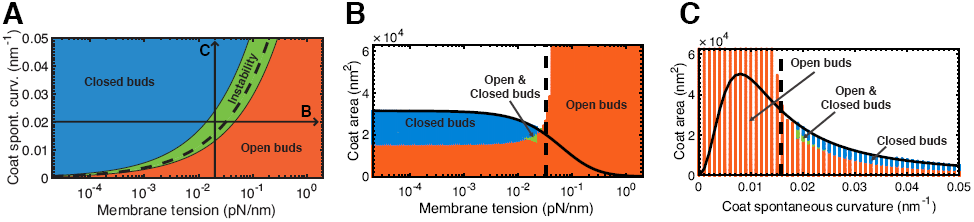
Bud morphology depends on bending rigidity, membrane tension, spontaneous curvature, and coat area. **(A)** Coat spontaneous curvature (*C*_0_) vs. membrane tension (λ_0_) phase diagram. The regions of the diagram are color coded according to the final shape of the membrane for coat “growing” simulations performed with the specified values for edge membrane tension and coat spontaneous curvature. Blue denotes closed, Ω-buds, Red denotes open, U-shaped pits, and Green are situations in which closed buds are obtained via a snapthrough transition. The snapthrough solutions cluster about the dashed line, Ves = 1, which separates the “high” and “low” membrane tension regimes (see main text). The lines labeled **B** & **C**, respectively, indicate the phase diagrams at right. **(B)** Coat area vs. membrane tension phase diagram, *C*_0_ = 0.02 nm^−1^. Blue denotes closed buds, red denotes open buds and green denotes parameters which have both open and closed bud solutions. The dashed line, Ves = 1, marks the transition from “low” to “high” membrane tension. The solid line represents the theoretical area of a sphere that minimizes the Helfrich energy at the specified membrane tension (see SI Appendix, Section 3). **(C)** Coat area vs. spontaneous curvature phase diagram, λ_0_ = 0.02pN/nm. The dashed line, Ves = 1, marks the transition between spontaneous curvatures that are capable and incapable of overcoming the membrane tension to form a closed bud. The solid line represents the theoretical area of a sphere that minimizes the Helfrich energy at the specified spontaneous curvature (see SI Appendix, Section 3).

These results can be understood by comparing the spontaneous curvature of the coat to the membrane tension and bending rigidity by studying the dimensionless quantity, 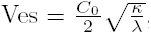, hereafter termed the *vesiculation number*. The dashed line in Figure 4A corresponds to Ves = 1, which bisects the low (Ves > 1) and high tension (Ves < 1) results. The snapthrough results cluster about this line, marking the transition region between the high and low tension cases. Importantly, this demonstrates that the preferred radius of curvature of the coat, 1/*C*_0_, must be smaller than the “natural” length scale of the membrane, 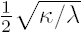 [27], for the coat to produce a closed bud in the absence of other mechanisms of curvature generation.

To study how the coat area affects the budding transition at a fixed spontaneous curvature, we varied spontaneous curvature and membrane tension for a fixed value of *C*_0_ = 0.02 nm^−1^ (Figure 4B). We also varied coat area against coat spontaneous curvature for a fixed value of λ_0_ = 0.02pN/nm (Figure 4C). For the sake of presentation, we here define Ω-shaped buds as any in which there is any overhang on the membrane contour (*ψ* > 90°, see SI Appendix, Figure S1), and U-shaped buds have no overhang (*ψ* <= 90°). Blue denotes Ω-shaped buds, the red region represents U-shaped buds, and green region indicates coexisting solutions of U- and *Ω*-shaped buds. In each plot, the dashed line represents Ves = 1, and the solid line is the area of a sphere with the given spontaneous curvature and membrane tension that would minimize the free energy (see SI Appendix, Section 3). We see that for Ves << 1 (high tension, low spontaneous curvature) the coat area can be increased arbitrarily high without formation of an *Ω*-shaped bud. Conversely, for Ves >> 1 (low tension, high spontaneous curvature), buds progress smoothly progress from U- to Ω-shaped buds as coat area is increased. Additionally, the final area of the coat before termination of the simulation closely aligns with the predicted area from energy minimization (see SI Appendix, Section 3).

### Increased coat rigidity smooths out the transition from open to closed buds

What properties of the membrane could be varied to overcome the instability at intermediate membrane tensions? Until now, we have taken the coat to have the same bending modulus as the bare membrane. The bending rigidity of clathrin-coated vesicles was estimated to be *κ*_CCV_ = 285 *k*_B_*T* = 2280 pN · nm from atomic force microscopy measurements [55]. Increasing the rigidity of the coated region to be *κ*_coat_ = 2400 pN · nm, 7.5 × the rigidity of the bare membrane *κ*_coat_ = 320 pN · nm, we conducted simulations at intermediate membrane tension (λ_0_ = 0.02pN/nm) with increasing coat area at constant spontaneous curvature (Figure 5A; Animation A6) and with increasing spontaneous curvature at constant area (Figure 5C; Animation A7). Comparing the plots of bud tip mean curvature as a function of coat area Figure (5B) and spontaneous curvature (Figure 5D), to those of the earlier simulations (Figures 3B and 3D, respectively), we see that there is now only a single branch of membrane shapes, indicating a smooth evolution from open, U-shaped buds to closed, Ω-shaped buds. We can understand these results by considering the vesiculation number. By increasing *κ*_coat_, we are increasing the value of the vesiculation number and are in effect shifting the phase space of bud morphologies toward the low tension regime.

**Figure 5.**
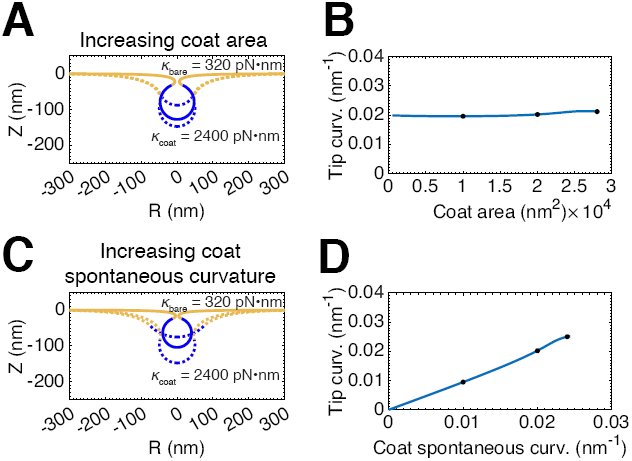
The snapthrough instability at physiological tension, λ_0_ = 0.02pN/nm, is abolished when the bending rigidity of the coat is increased relative to the bare membrane, *κ*_bare_ = 320 pN · nm, *κ*_coat_ = 2400 pN · nm. **(A)** Membrane profiles showing a smooth progression of bud morphologies as the area of the coat is increased (*A*_coat_ = 10, 000 nm^2^, 20,000 nm^2^, 28,000 nm^2^), *C*_0_ = 0.02 nm^−1^. **(B)** Mean curvature at the bud tip as a function of the area of the coat. The marked positions denote the membrane profiles shown in (A). There is now only a single branch of solutions (as compared to Figure 3B), indicating a smooth evolution from a flat membrane to a closed bud. **(C)** Membrane profiles showing a smooth progression of bud morphologies as spontaneous curvature of the coat is increased (*C*_0_ = 0.01 nm^−1^, 0.02 nm^−1^, 0.024nm^−1^), *A*_coat_ = 20, 106 nm^2^. (D) Mean curvature at the bud tip as a function of the spontaneous curvature of the coat showing a single branch of solutions (compare to Figure 3D).

### Force from actin polymerization can mediate the transition from a U-to Ω-shaped bud

What other mechanisms of force generation enable the cell to avoid the instability? Experiments have demonstrated that CME is severely affected by a combination of elevated tension and actin inhibition [5, 32]. To examine whether a force from actin polymerization is sufficient to induce a transition from open to closed bud morphologies, we modeled the force from actin polymerization in two orientations since the ultrastructure of the actin cytoskeleton at CME sites in live cells, and hence the exact orientation of the applied force, is currently unknown.

In the first candidate orientation, illustrated schematically in Figure 6A, actin polymerizes in a ring at the base of the pit with the network attached to the coat (via the actin-binding coat proteins Hip1R in mammals and its homologue Sla2 in yeast [56]). This geometric arrangement serves to redirect the typical compressive force from actin polymerization [57–59] into a net inward force on the bud and an outward force on the ring at the base of the invagination. This is analogous to the force from actin polymerization in yeast CME [31]. In the calculations, we take the force intensity to be homogeneously applied to the coated region, and the force intensity at the base is set such that the net applied force on the membrane integrates to zero. We find that an applied inward force of 15 pN on the bud is sufficient to drive the membrane from an open to closed configuration (Figure 6B; Animation A8, left). This force is well within the capability of a polymerizing branched actin network [60].

**Figure 6.**
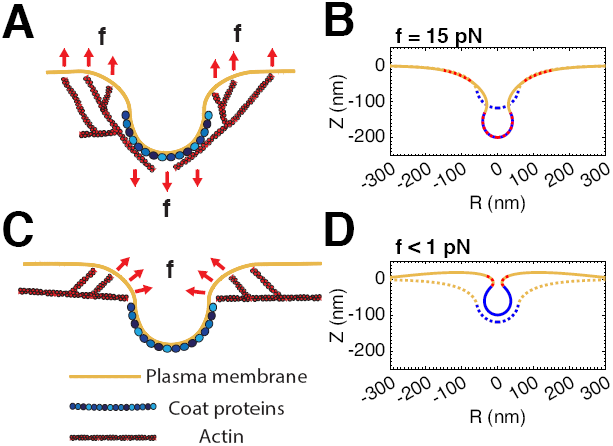
A force from actin assembly can mediate the transition from a U- to Ω-shaped bud, avoiding the instability at intermediate membrane tension, λ_0_ = 0.02pN/nm. Two orientations of the actin force were chosen based on experimental evidence from yeast [31] and mammalian [45] cells. **(A)** Schematic depicting actin polymerization in a ring at the base of the pit with the network attached to the coat, causing a net inward force on the bud. **(B)** At constant coat area, *A*_coat_ = 17, 593 nm^2^, and spontaneous curvature, *C*_0_ = 0.02 nm^−1^, a force (red dash) adjacent to the coat drives the shape transition from a U-shaped (dashed line) to Ω-shaped bud (solid line). The force intensity was homogeneously applied to the entire coat, and the force intensity at the base of the pit was set such that the total force on the membrane integrates to zero. The final applied inward force on the bud was **f** = 15pN, well within the capability of a polymerizing actin network [60]. **(C)** Schematic depicting actin assembly in a collar at the base, directly providing a constricting force. [45] **(D)** A constricting force (red dash) localized to the coat drives the shape transition from a U-shaped (dashed line) to Ω-shaped bud (solid line), *A*_coat_ = 17, 593 nm^2^, *C*_0_ = 0.02 nm^−1^. The force intensity was homogeneously applied perpendicular to the membrane to an area of 5, 027 nm^2^ immediately adjacent to the coated region. The final applied force on the membrane was **f** < 1pN.

In the second orientation, actin assembles in a collar at the base, directly providing a constricting force (Figure 6C), as suggested by the results of Collins et al. [45]. In the calculations, we take this force intensity to be oriented perpendicular to the membrane and applied homogeneously to a region immediately adjacent to the coat. This orientation produces a small vertical force on the membrane that is implicitly balanced by a force at the boundary of the domain through the boundary condition *Z* = 0nm. This counter force could easily be provided by the attachment of the underlying actin cortex to the plasma membrane [61]. Application of this constriction force is also sufficient to induce a smooth transition from U- to Ω-shaped buds with < 1pN of applied force (Figure 6D; Animation A8, right).

### A combination of increased coat rigidity and actin polymerization ensures robust vesiculation at high membrane tension

Since both increased coat rigidity and force from actin polymerization are sufficient to induce a smooth transition from open to closed buds at intermediate tension, we asked whether these prescriptions alone or in combination are sufficient to overcome high membrane tension to produce closed buds. In Figure 7A, we see that application of the inward directed force (as in Figure 6A) causes the membrane to deform from its initially flat morphology (dashed line) to a tubulated morphology (solid line). This final morphology is not especially reminiscent of CME, and the force required is now 60 pN which would require several dozen free actin filament plus ends, which may not be realistic for one endocytic site. Increasing the bending rigidity of the coat alone is also insufficient to produce a closed bud (Figure 7B, dashed line). However, this increased rigidity in combination with force from actin polymerization is sufficient to induce a transition to an Ω-shaped bud (solid line). Increasing membrane rigidity reduces the magnitude of the applied force by a factor of three to 20 pN, which is more biologically reasonable. In Figure 7C, we see that application of the constricting force (as in Figure 6C) is sufficient to induce a transition to a closed bud (solid line) from the initial flat morphology (dashed line). However, the magnitude of the applied force is now 160 pN, which is unrealistically high given the small area to which the force must be applied. Instead, by increasing the rigidity of the coat in combination with this actin force, we again obtain a closed bud (Figure 7D), but now the required force has been reduced by an order of magnitude to 16 pN.

**Figure 7.**
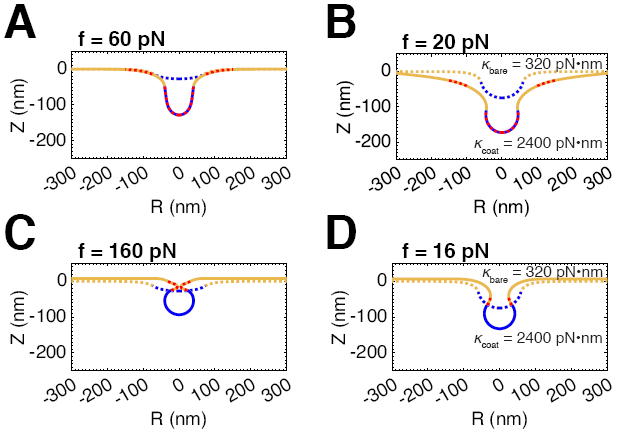
A combination of increased coat rigidity and force from actin polymerization ensures robust vesiculation, even at high membrane tension, λ_0_ = 0.2pN/nm, *C*_0_ = 0.02nm^−1^. **(A)** Application of the inward directed actin force (as in Figure 6A) induces tubulation, but not vesiculation, at high tension. **(B)** Increasing the stiffness of the coat alone is insufficient to overcome high membrane tension (dashed line). However, increasing the coat stiffness enables the applied force to induce vesiculation and decreases the magnitude of the force required by a factor of 3. **(C)** Application of the constricting actin force (as in Figure 6C) is sufficient to induce vesiculation, even at high tension. The magnitude of the applied force required is likely unrealistically high in a biologically relevant setting. **(D)** Increasing the coat stiffness decreases the force required to induce vesiculation by an order of magnitude.

## Discussion

In this study, we have investigated the role of membrane tension in governing the morphological landscape of CME and found that a combination of membrane tension and protein-induced spontaneous curvature governs the morphology of the endocytic pit (Figures 2). Additionally, we found that at intermediate membrane tensions, the bud must go through a snapthrough to go from an open to closed configuration (Figure 3). A key result from this work is that the vesiculation number can be used to identify the regimes of tension and curvature-mediated effects that separate the closed and open bud morphologies (Figure 4). Finally, we found that a modeling force generated by actin polymerization in CME can mediate the transition between open and closed buds at physiologically relevant membrane tensions (Figure 6). We believe that these results can explain and provide insight into the observations of a number of recent experimental studies.

**Figure 8.**
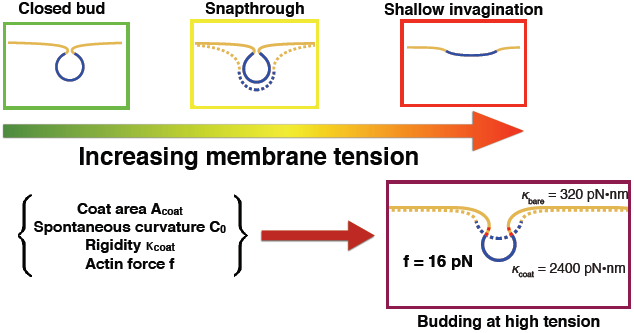
Design principles for robust vesiculation. The rigidity of the plasma membrane as well as the membrane tension resist budding by curvature-generating coats. In the low tension regime, as defined by the vesiculation number, increasing the coat area or spontaneous curvature is sufficient to induce a smooth evolution from a flat membrane to a closed bud. A combination of increased coat rigidity and force from actin polymerization is necessary to ensure robust vesiculation in the high membrane tension regime.

Boulant et al. [5] found that hypoosmotic shock and treatment with jasplakinolide (an actin stabilizing drug) has a severe inhibitory effect on CME in polarized MDCK and BSC-1 cells, while treatment with jasplakinolide alone did not affect CME dynamics. In light of our findings, it probable that the high tension induced by hypoosmotic shock resulted in a regime where the coat alone is insufficient to produce closed buds. The observed overabundance of U-shaped, presumably stalled, pits is consistent with a situation in which the membrane tension is in the snapthrough or high tension regime and coat assembly is unable to deform the membrane into a closed bud shape. Thus, under conditions of hypoosmotic shock it seems that a force exerted by the actin cytoskeleton, as in Figure 6, is necessary form a closed bud.

Saleem et al [22] used micropipette experiments to control the tension in the membrane of giant unilamellar vesicles (GUVs) to which they added purified coat components. We calculated the vesiculation number for the membrane tensions (≈ 0.5 – 3pN/nm) set by micropipette aspiration to be less than 1 over a wide range of spontaneous curvatures, indicating a high membrane tension regime in their set up. Thus, our model is consistent with their observations of shallow buds observed in isotonic conditions. One result that our model cannot explain is the lack of any clathrin assembly observed under hypotonic conditions. It is possible that at extremely high membrane tensions the coat is simply unable to stay bound to the membrane at the extremely flat morphology that would be expected.

Avinoam et al. found that the size of the coat does not change substantially during membrane deformation in CME [46]. This is in contrast to the canonical view that the clathrin coat should directly impose its preferred curvature on the underlying membrane [8]. There are two possible explanations for this observation in the context of our study. One is that the membrane tension is too high for the coat to deform the membrane, so that other mechanisms of curvature generation (e.g. actin polymerization or BAR domain-induced curvature) are necessary to remodel the membrane. The second is that the coat undergoes a “maturation” process that progressively increases its spontaneous curvature, and hence its capability to bend the membrane, as in Figure 2B. The observation that actin inhibition causes substantial defects in CME in this cell type [32] is consistent with the hypothesis that the membrane tension could be elevated in this cell type, though this would need to be confirmed experimentally. Thus, it is possible that the observation that the size of the clathrin coat is constant during the budding process might be specific to SK-MEL-2 cells and in particular on the typical membrane tension of this cell line.

Our results also build on previous models that have been used to study CME. We have shown here that membrane deformation at high tension can be achieved by coupling increased coat rigidity and actin-mediated forces (Figure 7). Walani et al. also explored budding at high tension and predicted that an actin-force-driven snapthrough instability could drive scission in yeast CME [42]. However, this instability is a consequence of the exact implementation of the actin force (SI Appendix, Figure S7; Animation A9), and so its physiological relevance is unclear.

Other models have assumed that the proteins exert a spherical cap and a line tension to form a bud [22, 39, 62]. Here we obtain buds as a result of protein-induced spontaneous curvature where the final radius of the bud depends on the membrane tension (Figure 4B; SI Appendix, Section 3). Line tension was not explicitly accounted for in our model since we used a smooth function to model the interface representing the heterogeneity of the membrane (SI Appendix, Figure S2). Line tension captures the energy of an interface, but by smoothing out this interface to a continuum with a sharp transition we are able to construct a single model for multiple domains.

Another aspect of heterogeneous membrane properties that we explored was variation in the Gaussian modulus between the coated and bare membrane, which has been demonstrated both theoretically [63] and experimentally [38] to affect the location of the phase boundary in the neck of phase-separated vesicles. In addition to affecting the location of the boundary relative to the neck, we found that variation in the Gaussian modulus has a profound effect on the progression of budding. Increasing the Gaussian modulus of the coat relative to the bare membrane inhibits budding, while decreasing it can smooth out the instability at intermediate membrane tension (SI Appendix, Figure S8). While interesting, until more is known about how the lipid and protein composition at endocytic sites affects the Gaussian modulus, it is unclear what relevance these results have in CME.

One aspect of CME not explicitly addressed by this study is that the endocytic machinery includes curvature-generating proteins outside of the coat proteins and the actin machinery. In particular, recent modeling studies have demonstrated that cyclindrical curvature induced by BAR-domain proteins can play an important role in reducing the force requirement for productive CME in yeast [27, 42]. However, CME is still productive in 50% of events even with complete knockout of the endocytic BAR-domain proteins in this organism [64], while actin assembly is absolutely required [25, 26]. Additionally, in mammalian cells a large percentage of CCPs were found to stall at high membrane tension when actin is inhibited [5] despite the fact that the BAR-domain proteins were presumably unaffected. These results suggest that while curvature generated by BAR-domain proteins may help to facilitate productive CME, force from actin assembly seems to be most important in challenging mechanical environments.

## Model predictions

Our model makes several experimentally testable predictions.
- There is conflicting evidence as to whether actin is an essential component of the endocytic machinery in mammalian cells [5, 21, 32]. We predict that CME in cell types with higher membrane tensions (i.e. Ves < 1) will be sensitive to perturbations to actin dynamics. Similarly, a reduction in membrane tension might relieve the necessity for actin polymerization in cells types where it has been found to be important for productive CME. A systematic study of the membrane tension in different cell types along with the sensitivity of CME to actin inhibitors will provide a strong test of the model and potentially clarify the role of actin in CME in mammalian cells.
- Reduction in the spontaneous curvature of the clathrin coat will have severe effects on CME dynamics at elevated membrane tension. A recent study by Miller et al. showed that depletion of the endocytic adaptor proteins AP2 and CALM resulted in smaller and larger CCPs, respectively [12]. This effect was attributed to the presence of a curvature-driving amphipathic helix in CALM and the fact that AP2 typically recruits bulkier cargos than CALM, which translates in our framework into a reduction of the coat spontaneous curvature upon CALM depletion. We predict that CME in cells depleted of CALM will be more sensitive to increase in membrane tension (and/or actin inhibition) than in cells depleted of AP2 because successful budding is predicted to be a function of both membrane tension and spontaneous curvature (Figure 4).
- Reduction in the stiffness of the coat will inhibit its ability to bend membranes, especially at elevated membrane tension. This effect could be directly tested in a reconstitution system similar to that of Saleem et al. [22] in the presence or absence of clathrin light chains which have been shown to modulate the stiffness of the clathrin lattice [65].

## Limitations of the model

Despite the agreement with experimental data and generation of model predictions, we acknowledge some limitations of our model. Our model is valid only for large length scale deformations, since the Helfrich energy is valid over length scales much larger than the thickness of the bilayer [13]. Further, we have assumed mechanical equilibrium for the membrane and future efforts will focus on including dynamics of the membrane. And finally, spontaneous curvature is one term that gathers many aspects of membrane bending while ignoring exact molecular mechanisms (protein insertion into the bilayer versus crowding). While it is effective for representing the energy changes to the membrane due to protein interaction, detailed models will be needed to explicitly capture the different mechanisms.

## Conclusions

Reductionist approaches in cell biology, while very powerful in identifying univariate behavior, can be limited in their conclusions because processes like CME are controlled by multiple variables. Using a “systems” approach, we have investigated a multivariate framework that identifies the fundamental design principles of budding. Despite the inherent complexities of protein-induced budding, we found that coat area, coat spontaneous curvature, bending moduli, and actin-mediated forces are general factors that can contribute to robust vesiculation against opposing forces like membrane tension.

Though we have primarily focused on budding in the context CME, our findings are general to any budding process. For example, it has been shown that membrane deformation by COPI coats is also inhibited by membrane tension [36] and that rigidity of the COPII coat is essential for export of bulky cargos [66]. Since the membranes of the endoplasmic reticulum and the Golgi are also under tension [67], we expect that the shape evolution of buds from these organelles is also determined by a balance of the coat spontaneous curvature, bending rigidity and membrane tension. Other membrane invaginations are also presumably governed by a similar set of physical parameters. For example, caveolae have been proposed to act as a membrane reservoir that buffers changes in membrane tension by disassembling upon an increase in membrane tension [23]. A similar framework to the one used in this study might provide some insight into the morphology and energetics of membrane buffering by caveolae. Moving forward, more detailed measurements of both the membrane tension within cells and the spontaneous curvature of various membrane-bending proteins will be essential to verify and extend the results presented here.

**Table 1:** Notation used in the model

| Notation | Description | Units |
| --- | --- | --- |
| $A_{\text{coat}}$ | Area covered by the coat | $\text{nm}^2$ |
| $C$ | Spontaneous curvature | $\text{nm}^{-1}$ |
| $\theta^\alpha$ | Parameters describing the surface | |
| $W$ | Local energy per unit area | $\text{pN}/\text{nm}$ |
| $\mathbf{r}$ | Position vector | |
| $\mathbf{n}$ | Normal to the membrane surface | unit vector |
| $\mathbf{a}_\alpha$ | Basis vectors describing the tangent plane, $\alpha \in \{1, 2\}$ | |
| $\lambda$ | Membrane tension, $-(W + \gamma)$ | $\text{pN}/\text{nm}$ |
| $p$ | Pressure difference across the membrane | $\text{pN}/\text{nm}^2$ |
| $H$ | Mean curvature of the membrane | $\text{nm}^{-1}$ |
| $K$ | Gaussian curvature of the membrane | $\text{nm}^{-2}$ |
| $\kappa$ | Bending modulus (rigidity) | $\text{pN} \cdot \text{nm}$ |
| $\kappa_G$ | Gaussian modulus | $\text{pN} \cdot \text{nm}$ |
| $k_B T$ | Units of thermal energy, $\approx 2.5 \text{ kJ/mol}$ | |

**Table 2:**
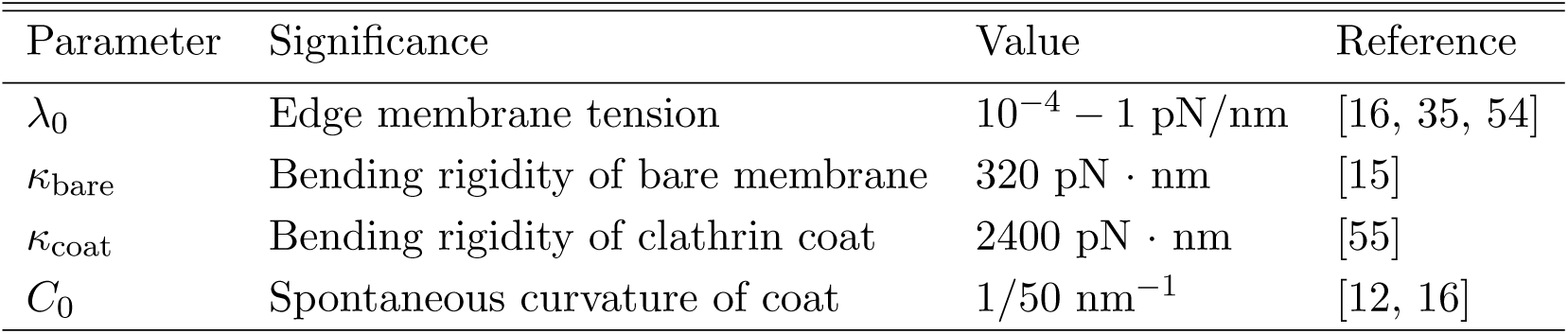
Parameters used in the model

| Parameter | Significance | Value | Reference |
| --- | --- | --- | --- |
| $\lambda_0$ | Edge membrane tension | $10^{-4} - 1 \text{ pN}/\text{nm}$ | [16, 35, 54] |
| $\kappa_{\text{bare}}$ | Bending rigidity of bare membrane | $320 \text{ pN} \cdot \text{nm}$ | [15] |
| $\kappa_{\text{coat}}$ | Bending rigidity of clathrin coat | $2400 \text{ pN} \cdot \text{nm}$ | [55] |
| $C_0$ | Spontaneous curvature of coat | $1/50 \text{ nm}^{-1}$ | [12, 16] |

## Acknowledgements

The authors would like to thank Matt Akamatsu, Charlotte Kaplan, and our anonymous reviewers for critical reading of the manuscript. This research was conducted with Government support under and awarded by DoD, Air Force Office of Scientific Research, National Defense Science and Engineering Graduate (NDSEG) Fellowship, 32 CFR 168a to J.E.H., the National Institutes of Health Grant R01GM104979 to G.O., the National Institutes of Health Grant R35GM118149 to D.G.D., and the UC Berkeley Chancellor's Postdoctoral Fellowship, the Air Force Office of Scientific Research award number FA9550-15-1-0124 and NSF PHY-1505017 to P.R.

References

